# Genetic diversity assessment of neem (*Azadirachta indica* A. Juss) in northern Nigeria

**DOI:** 10.1101/2021.11.22.469531

**Authors:** Barka P. Mshelmbula, Geoffrey O. Anoliefo, Beckley Ikhajiagbe, Boniface O. Edegbai

**Affiliations:** Department of Plant Science and Biotechnology, Federal University of Lafia, PMB 146, Lafia, Nigeria; Environmental Biotechnology and Sustainability Research Group, Dept. of Plant Biology and Biotechnology, University of Benin, Benin City, Nigeria; Applied Environmental Biosciences and Public Health Research Group, Department of Microbiology, University of Benin, Benin City, Nigeria

**Keywords:** neem, molecular characterization, heritability, genetic gain, *Azadirachtaindica*

## Abstract

Neem is a tropical tree that can adapt to a wide range of places and particularly to semi-arid conditions. As at present, it is grown in many Asian countries and also in the tropical regions of the western hemisphere. Genetic variability and diversity are a major requirement needed for both immediate results and the one ones thereafter adaptation of plant types in their original domain. The evaluation of genetic diversity of any species is extremely crucial for their sustainability, continuity, survival and gene manipulation. Major breakthroughs in the field of molecular biology was able to develop several tools for the investigation of genetic diversity at the genome level to determine phylogenetic relationships among inter or intra-species. The advent of molecular markers for the detection and exploitation of DNA polymorphism is one of the major breakthroughs in the world of molecular genetics.

The importance of genetic diversity in plant germplasm conservation, especially in economically important species such as *Azadirachtaindica*, is enormous, particularly in Nigeria. The question is whether *A. indica* from different Agro-ecological zones have genetic variations or similarities. This was the bane of the current study, which used RAPD to look atgenetic diversity of 27 randomly selected neem trees within the agro-ecological zones in Northern Nigeria. A total of 9 primers were employed out of which only 5 were responsive (OPA-02, OPA-03, OPA-15 and OPA-19). These primers showed dissimilarities in the visible DNA bands among the various tree samples. There was evidence of genetic dissimilarities among the trees sampled. Differences in percentage polymorphism was reported, where it was reportedly highest among the Borno State tree samples (97.44%), compared to those in Yobe State with no polymorphism.

## 1. Introduction

*Azadirachtaindica* A. Juss (Meliaceae) is a well adaptable plant that lives in forests where agriculture is practiced of the semiarid and arid tropics, and is native to the dry forest zones of Asia (Koul*et al*. 1990, Ketkar and Ketkar 1995). It is widely distributed, and this may be the result of its wide acceptability, particularly for its economic importance.Neem trees have been broadly developed within the tropics. Its seedlings are raised in the nursery from where they are transplanted unto the field. It is a tree in Northern Nigeria that is almost synonymous with the term “tree of life”. This is so because there is virtually hardly any man-made settlement without the neem tree. It flourishes in nutrient-poor dry soils and is tolerant of temperatures when high but is vulnerable to over the top cold or ice. It has uses in ethno medicine, construction, energy and fuel, environmental protection, traditional significance, fencing and agricultural intervention. Neem trees have been broadly developed within the tropics. Its seedlings are often grown in the nursery and later raised in the field.

Due to the cosmopolitan nature of the tree, it may be exposed to a wide range of environments and ecosystems, particularly since it has been connected to nearly every agro-ecological zone in Nigeria.It’s unclear if this ability is inherited or provides genetic diversification. Genetic diversity is an essential factor in species stability in any ecosystem, as it provides important transformations to the dominant biotic and abiotic natural conditions, as well as allowing changes in genetic make-up to adapt to changes in the environment.

The assessment of hereditary difference of any species is extremely crucial for their preservation and gene improvement (Khan et al. 2012). The objective of this study is to evaluate the genetic variability among individual neem species using Randomly Amplified Polymorphic DNA (RAPD). The study also hopes to determine the genetic similarities and differences among the individuals of the selected plant population within the agro ecological zones of northern Nigeria.Dhillion et al. (2007) utilized RAPD markers to assess the genetic variety in *Azadirachtaindica* population from various eco-geological locale of India.

## 2. Materials and Methods

### 2.1 Plant Sample Collection

The study occurred between February and July of 2019. The research randomly targeted 27 individual Neem tree species scattered within the 4 agro-ecological zones of Northern Nigeria (Figure 1). The plant samples were not obtained from the wild; they were obtained from within cultivated or built environment. This is because the plant is very common either as wind break, aesthetic, horticultural or for phytomedicinal purposes. Formal identification of the samples was made possible with assistance from Plant Taxonomy and Herbarium Unit of the Department of Plant Biology and Biotechnology, University of Benin, Benin City, and the Department of Botany, Federal University, Lafia, Nassarawa State, Nigeria. Care was taken to ensure that tree species of interest were not sampled within a single, but a wider location in so far as they represent designated ecological zones of interest.

**Figure 1:**
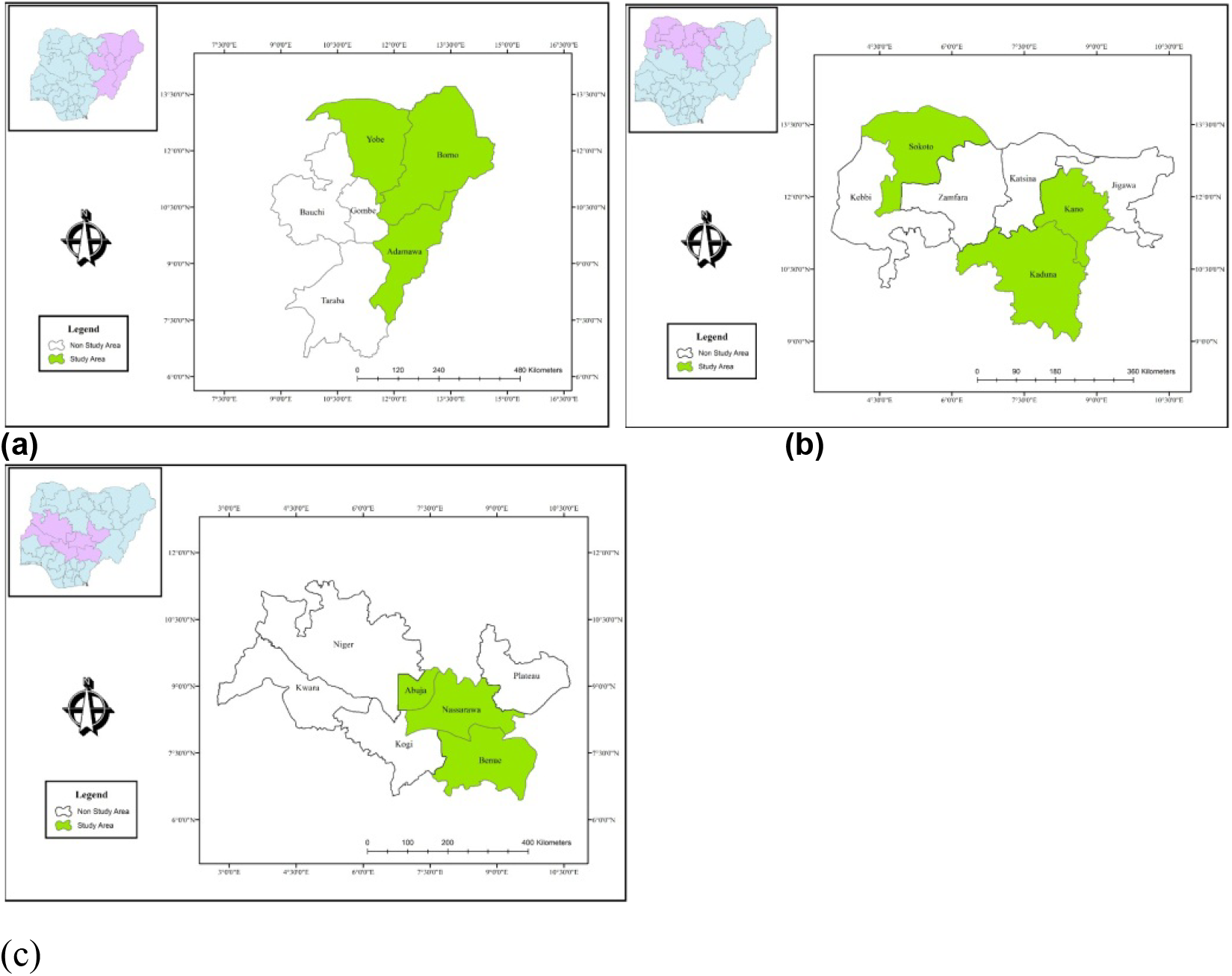
Map showing study area at the (a) North Eastern, (b) North West and the (c) North Central region of the country

### 2.2 Sample collection and storage

For RAPD analysis, seeds were used. Seeds from mature fruits were collected from each tree as described by Singh *et al. (*2005). Selection of tree for sample collection was based on the following conditions; that the tree should have a visible bole of not less than 3m, a basal girth of not less than 50 cm and must show evidence of fruiting. Seed samples were immediately covered with aluminium foil paper and placed in ice packed coolers. The seeds were then labeled according to geographic locations of trees as provided on Table 1.

**Table 1:** Sample codes for collection of plant samples during the study.

| Code | Plant physical location | State | Agro-ecological zone |
| --- | --- | --- | --- |
| YB-BK-1 | Along Malari motor park, Malari ward, Damaturu. | Yobe | Sahel savanna |
| YB-BK-2 | Potiskum cattle market | Yobe | Sahel savanna |
| YB-BK-3 | Federal University, Gashua. Sabongari, Nguru-Gashua Road, Damasak road, Gashua | Yobe | Sahel savanna |
| AD-BK-1 | Federal college of education, Yola | Adamawa | Southern Guinea savanna |
| AD-BK-2 | Emirs palace, Mubi | Adamawa | Southern Guinea savanna |
| AD-BK-3 | College of Agriculture, Ganye | Adamawa | Southern Guinea savanna |
| BN-BK-1 | E.Y.N Church, Wulari, Maiduguri | Borno | Sahel savanna |
| BN-BK-2 | Army University, Biu | Borno | Sahel savanna |
| BN-BK-3 | University of Maiduguri | Borno | Sahel savanna |
| KN-BK-1 | GidanBuhari Road, T/Wada Kano | Kano | Sudan savanna |
| KN-BK-2 | BUK, Kano state | Kano | Sudan savanna |
| KN-BK-3 | Dala Orthopedic Hospital, Kano | Kano | Sudan savanna |
| SK-BK-1 | Usman Dan Fodio University, Sokoto | Sokoto | Sudan savanna |
| SK-BK-2 | SaniDingyadi Sec. Sch. Farfaru, Sokoto | Sokoto | Sudan savanna |
| SK-BK-3 | Sokoto main market, Sokoto | Sokoto | Sudan savanna |
| KD-BK-1 | Ahmadu Bello University, Zaria | Kaduna | Northern Guinea savanna |
| KD-BK-2 | Sabo motor park, Kaduna | Kaduna | Northern Guinea savanna |
| KD-BK-3 | Close to Stanbic IBTC | Kaduna | Northern Guinea savanna |
| NS-BK-1 | OBKi-1 opposite Nasarawa Polytechnic, Lafia | Nasarawa | Southern Guinea savanna |
| NS-BK-2 | Hope academy secondary school, Akwanga | Nasarawa | Southern Guinea savanna |
| NS-BK-3 | Awe Motor Park, Awe LGA | Nasarawa | Southern Guinea savanna |
| AJ-BK-1 | Lungi Barracks, Maitama Abuja. | Abuja | Derived savanna |
| AJ-BK-2 | MaBKilla Barracks, Asokoro Abuja | Abuja | Derived savanna |
| AJ-BK-3 | Opposite State House Medical Centre, Asokoro Abuja | Abuja | Derived savanna |
| BN-BK-1 | University of Agriculture, Makurdi | Benue | Derived savanna |
| BN-BK-2 | Benue State University, Makurdi | Benue | Derived savanna |
| BN-BK-3 | Peace House, Gboko | Benue | Derived savanna |

### 2.3 RAPD Analysis

In order to obtain clear reproducible amplification products, different preliminary experiments were carried out in which a number of factors were optimized. These factors included PCR annealing temperatures and concentration of each of the template DNA (Kumari and Thakur, 2014). A total of 9 random DNA oligonucleotides were independently used in the PCR reaction (Table 2). PCR was performed with 10 μl volumes using Thermo Fisher Scientific PCR mix. The final concentrations were as follows: 1 X Taq buffer, 2.5 mM, MgCl_2_, 0.32 mM, dNTP mixture, 0.25 μM for each primer, 0.5 U Taq polymerase and 10 ng genomic DNA. All reactions were performed using a Wagetech Projects Master Cycler (Eppendorf, Hamburg, Germany) with an initial denaturing cycle of 5min at 95 ^°^C, 40 cycles of 30 s at 93 ^°^C, 1min at 43 ^°^C, 1 min at 72 ^°^C, and a final extension cycle of 10 min at 72 ^°^C. The pure DNA was stored at – 20 ^°^C.

**Table 2:**
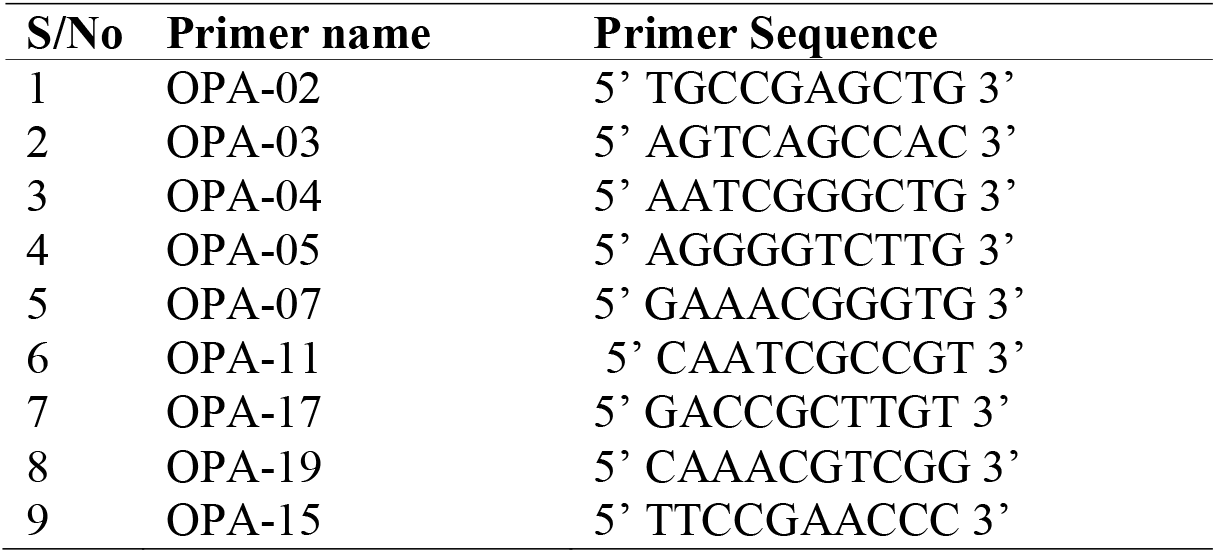
Primer information on 9 primers for RAPD study.

### 2.4 Agarose Gel Electrophoresis

The PCR products were visualized using ethidium bromide dye in 2 % agarose gels. The gel matrix was prepared following the protocol of Ihase et al. (2014). The solution was then poured on a gel tray with a comb inserted in it. Using a micro pipette, 1 µl of loading dye was added to 9 µl of the final product and the mixture was loaded into the wells. The tank was connected to a power source at 90 V for 60 mins. The gel photograph was taken using a digital camera.

### Data Analysis

The DNA bands were counted and fragment sizes compared with those of the DNA Ladder. The bands were scored and used for genotyping. The presence or absence of each DNA band was treated as a binary character in a data matrix (coded 1 and 0, respectively) to calculate genetic similarity and to predict variations among the fruit forms. The genetic distance between the three fruit forms was calculated using GeneAlEx 6.502 software which was used to construct a dendogram on MEGA 7.0.21 software (Kumar et al. 2016).

## 3.0 RESULTS

Nine (09) random primers were obtained and screened for responsiveness to the neem genome. This was done to select the most suitable primer for the RAPD study. Only primers OPA-02, OPA-03, OPA-15 and OPA-19 turned out to be responsive. Primers OPA-04, OPA-05, OPA 6, and OPA-11 were not responsive. They did not amplify any region of the neem genome tested. Information of the random primers used for this study are presented in Table 2. RAPD primers variations in the study of genetic diversity of Neem tree has been presented on Table 3. Among the primers used in the study, Primer OPA-15 was most effective. Mean number of effective alleles ranged from 1.095 – 1.625. OPA-15 had the highest diversity index compared to others used in the study. Expected Heterozygosity ranged from 0.088 – 0.405. heterozygocity was higher with Primer OPA-15.

**Table 2:** Random primer screening.

| **S/No** | **Primer name** | **Primer Sequence** | **Response** |
| --- | --- | --- | --- |
| 1 | OPA-02 | 5' TGCCGAGCTG 3' | + |
| 2 | OPA-03 | 5' AGTCAGCCAC 3' | + |
| 3 | OPA-04 | 5' AATCGGGCTG 3' | + |
| 4 | OPA-05 | 5' AGGGGTCTTG 3' | No response |
| 5 | OPA-07 | 5' GAAACGGGTG 3' | - |
| 6 | OPA-11 | 5' CAATCGCCGT 3' | No response |
| 7 | OPA-17 | 5' GACCGCTTGT 3' | - |
| 8 | OPA-19 | 5' CAAACGTCGG 3' | + |
| 9 | OPA-15 | 5' TTCCGAACCC 3' | + |

**Table 3:** RAPD primers variations in the study of genetic diversity of Neem tree.

| Primers | Na | Ne | I | He | uHe |
| --- | --- | --- | --- | --- | --- |
| OPA-04- 1 | 0.444 | 1.153 | 0.129 | 0.088 | 0.105 |
| OPA-04- 2 | 0.556 | 1.095 | 0.106 | 0.067 | 0.080 |
| OPA-04- 3 | 0.556 | 1.212 | 0.151 | 0.108 | 0.130 |
| OPA-04- 4 | 1.000 | 1.307 | 0.257 | 0.175 | 0.210 |
| OPA-04- 5 | 1.000 | 1.365 | 0.280 | 0.196 | 0.235 |
| OPA-04- 6 | 1.000 | 1.307 | 0.257 | 0.175 | 0.210 |
| OPA-04- 7 | 1.000 | 1.307 | 0.257 | 0.175 | 0.210 |
| OPA-04- 8 | 0.667 | 1.201 | 0.182 | 0.121 | 0.145 |
| OPA-04- 9 | 0.889 | 1.190 | 0.212 | 0.133 | 0.160 |
| OPA-02-1 | 0.667 | 1.259 | 0.204 | 0.142 | 0.170 |
| OPA-02-2 | 0.667 | 1.259 | 0.204 | 0.142 | 0.170 |
| OPA-02-3 | 0.667 | 1.259 | 0.204 | 0.142 | 0.170 |
| OPA-02-4 | 0.667 | 1.259 | 0.204 | 0.142 | 0.170 |
| OPA-02-5 | 0.667 | 1.259 | 0.204 | 0.142 | 0.170 |
| OPA-02-6 | 0.556 | 1.153 | 0.129 | 0.088 | 0.105 |
| OPA-02-7 | 0.556 | 1.153 | 0.129 | 0.088 | 0.105 |
| OPA-02-8 | 0.778 | 1.201 | 0.182 | 0.121 | 0.145 |
| OPA-02-9 | 0.889 | 1.249 | 0.235 | 0.154 | 0.185 |
| OPA-02-10 | 0.444 | 1.153 | 0.129 | 0.088 | 0.105 |
| OPA-03-1 | 0.778 | 1.259 | 0.204 | 0.142 | 0.170 |
| OPA-03-2 | 0.889 | 1.259 | 0.204 | 0.142 | 0.170 |
| OPA-03-3 | 0.889 | 1.201 | 0.182 | 0.121 | 0.145 |
| OPA-03-4 | 0.889 | 1.201 | 0.182 | 0.121 | 0.145 |
| OPA-03-5 | 1.000 | 1.307 | 0.257 | 0.175 | 0.210 |
| OPA-03-6 | 1.000 | 1.249 | 0.235 | 0.154 | 0.185 |
| OPA-03-7 | 1.000 | 1.307 | 0.257 | 0.175 | 0.210 |
| OPA-03-8 | 0.778 | 1.201 | 0.182 | 0.121 | 0.145 |
| OPA-15-1 | 0.667 | 1.201 | 0.182 | 0.121 | 0.145 |
| OPA-15-2 | 0.667 | 1.201 | 0.182 | 0.121 | 0.145 |
| OPA-15-3 | 0.889 | 1.249 | 0.235 | 0.154 | 0.185 |
| OPA-15-4 | 1.333 | 1.402 | 0.363 | 0.242 | 0.290 |
| OPA-15-5 | 1.778 | 1.614 | 0.515 | 0.350 | 0.420 |
| OPA-15-6 | 1.667 | 1.566 | 0.462 | 0.317 | 0.380 |
| OPA-15-7 | 1.667 | 1.625 | 0.484 | 0.338 | 0.405 |
| OPA-19-1 | 0.778 | 1.201 | 0.182 | 0.121 | 0.145 |
| OPA-19-2 | 0.889 | 1.259 | 0.204 | 0.142 | 0.170 |
| OPA-19-3 | 0.889 | 1.259 | 0.204 | 0.142 | 0.170 |
| OPA-19-4 | 1.000 | 1.307 | 0.257 | 0.175 | 0.210 |
| OPA-19-5 | 1.000 | 1.307 | 0.257 | 0.175 | 0.210 |
| Grand mean | 0.875 | 1.270 | 0.228 | 0.155 | 0.186 |

*Na* = No. of different alleles; *Ne* = No. of effective alleles; *I* = Shannon's Information Index; *He* = Expected Heterozygosity; *uHe* = Unbiased Expected Heterozygosity

### Preliminary gels from responsive primers

Using Primer OPA-02, visible bands were reported in samples collected from Abuja-2, Adamawa-3, Benue-1, Benue-2, Borno-3, Sokoto-3 and Yobe-2 (Figure 2). The implication is that among the samples collected from Abuja, one was genetically distinct from the other. This was similar to those collected in Adamawa, Borno, Benue, Sokoto and Yobe States respectively.Using Primer OPA-03 (Figure 3), amplifications were recorded for samples collected from Abuja-1, Abuja-2, Abuja-3, Kaduna-1,. Kaduna-2, Borno-3, Nassarawa-1, Nassarawa2, Yobe-1, Yobe-2, Yobe-3. Using Primer OPA-02, samples collected from Yobe State were of similar genetic identity as their banding patterns were similar. They also had similar amplifications with samples collected from Borno State.

**Figure 2:**
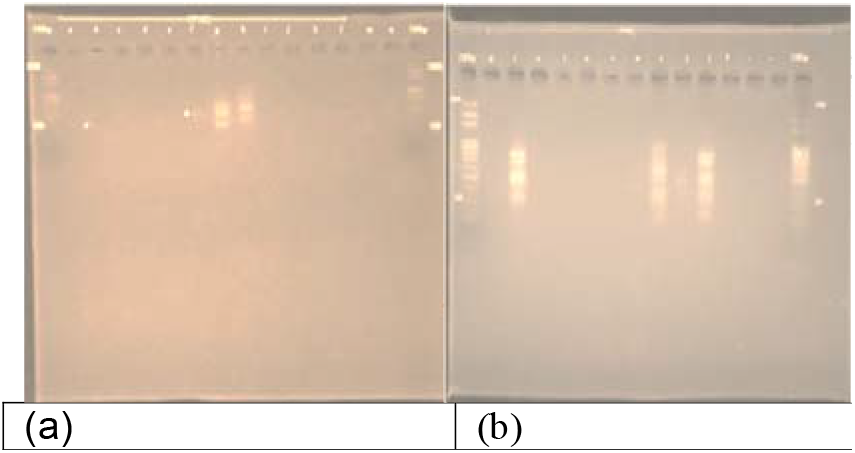
Gels of responsive Primer OPA-02

**Figure 3:**
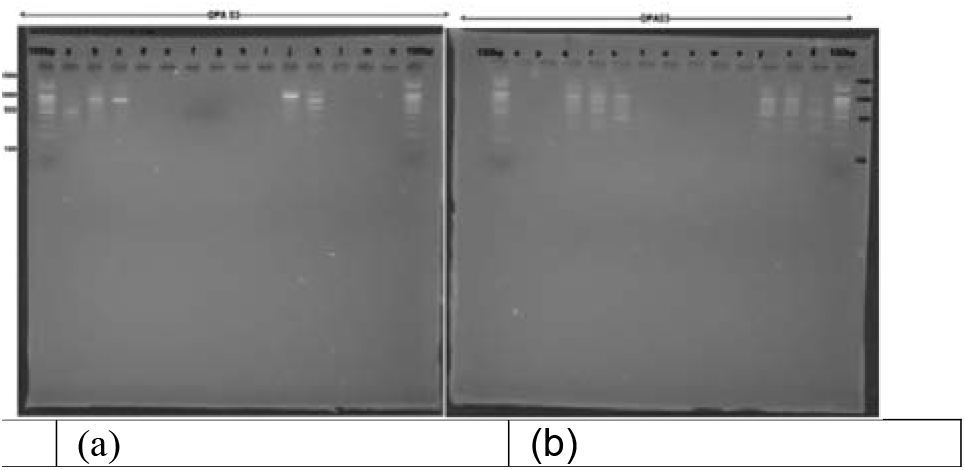
Gels of responsive Primer OPA-03

Gels of responsive Primer OPA-04 showed amplifications for all Abuja samples, Kaduna-1 and Kaduna-2, Borno-2 and Borno-3, Nasarawa-1, as well as all samples from Yobe State (Figure 4). These identities had distinct banding patterns indicating genetic dissimilarities. More than 80% of the samples collected showed amplified bands using Primer OPA-15, with majorly dissimilar characteristics (Figure 5). Samples from Borno and Yobe States had similar banding patterns, indicating possible genetic identities (Figure 6).

**Figure 4:**
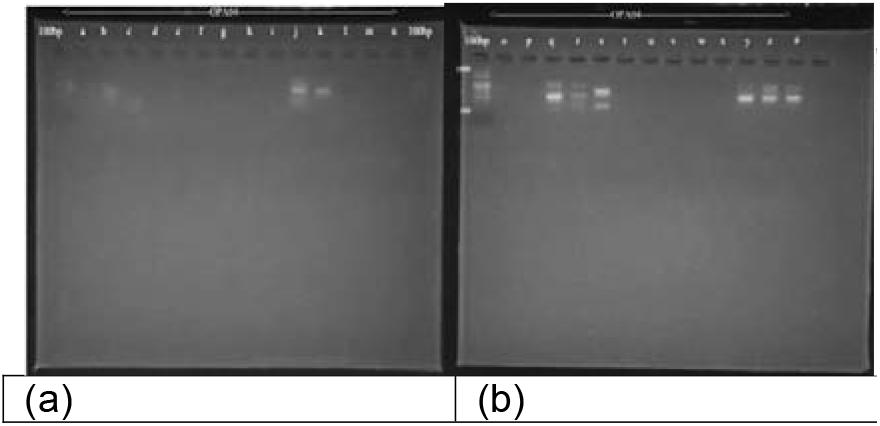
Gels of responsive Primer OPA-04

**Figure 5:**
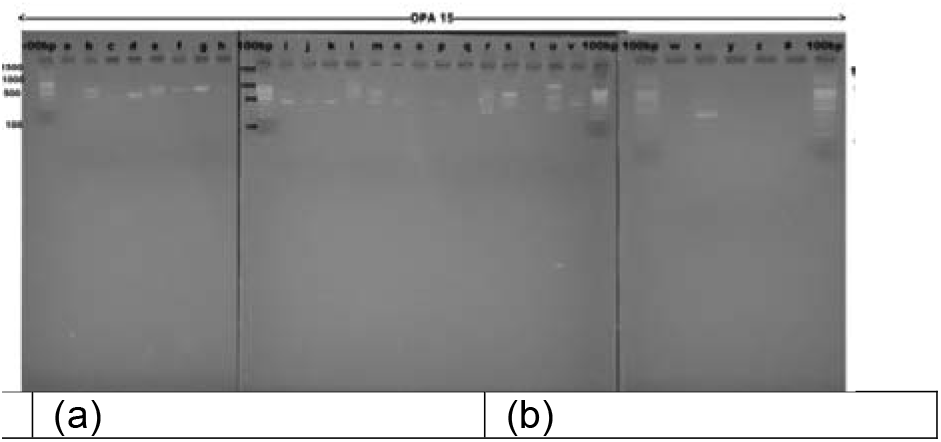
Gels of responsive Primer OPA-15

**Figure 6:**
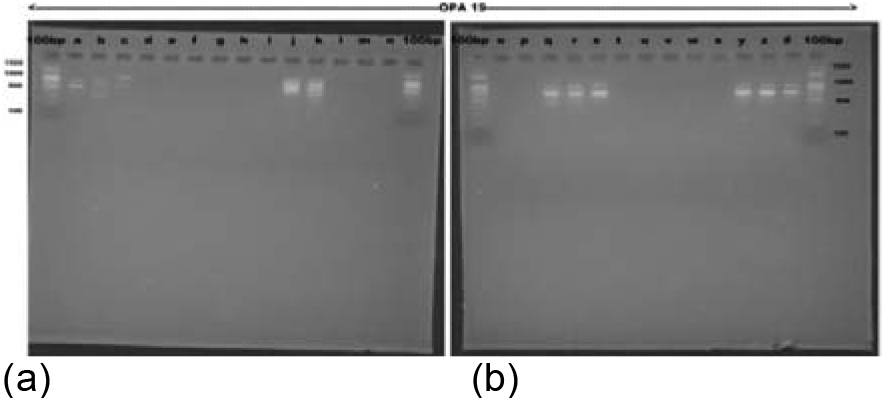
Gels of responsive Primer OPA-19 Abuja 2-b, Abuja 3-c; Adamawa 1-d, Adamawa 2-e, Adamawa 3-f; Benue 1-g, Benue 2-h, Benue 3-I; Kaduna 1-j, Kaduna 2-k, Kaduna 3-L; Kano 1-m, Kano 2-n, Kano 3-O; Borno 1-p, Borno 2-q, Borno 3-r; Nasarawa 1-s, Nassarawa 2-t, Nassarawa 3-u; Sokoto 1-v, Sokoto-2; w, Sokoto 3 x; Yobe 1-y, Yobe 2-z, Yobe 3-#

In the Abuja samples, 20 bands were observed; all these Different Bands had a Frequency ≥ 5% (Table 4). However, the Borno samples had the highest number of bands (38). Of the 38 bands in the Borno samples, 20 were locally common bands found in 50% or fewer populations. Mean number of available alleles ranged from 0.154 (Kano samples) to 1.949 (Borno samples) (Table 5). In spite of this difference, number of effective alleles between these two locations were comparable (1.046 – 1.686).Table 6 shows the percentage polymorphism of RAPD markers across the nine locations in the study. Percentage polymorphism was highest among the Borno samples (97.44%), compared to samples in Yobe State with no polymorphism. Mean percentage polymorphism was 39.60%.

**Table 4:** Amplified bands detected with thirty-nine primers in Neem tree species across nine states in Nigeria (Primer OPA-15)

| Population | Abuja | Adamawa | Benue | Kaduna | Kano | Borno | Nasarawa | Sokoto | Yobe |
| --- | --- | --- | --- | --- | --- | --- | --- | --- | --- |
| No. Bands | 20 | 6 | 13 | 29 | 3 | 38 | 25 | 15 | 19 |
| No. Bands Freq. $\geq 5\%$ | 20 | 6 | 13 | 29 | 3 | 38 | 25 | 15 | 19 |
| No. Private Bands | 0 | 0 | 0 | 0 | 0 | 0 | 0 | 0 | 0 |
| No. LComm Bands ($\leq 25\%$) | 0 | 0 | 0 | 1 | 0 | 2 | 0 | 1 | 0 |
| No. LComm Bands ($\leq 50\%$) | 3 | 2 | 9 | 11 | 0 | 20 | 7 | 11 | 5 |
| Mean He | 0.144 | 0.061 | 0.143 | 0.261 | 0.028 | 0.389 | 0.221 | 0.145 | 0.000 |
| SE of Mean He | 0.031 | 0.024 | 0.034 | 0.031 | 0.016 | 0.018 | 0.029 | 0.036 | 0.000 |
| Mean uHe | 0.173 | 0.073 | 0.172 | 0.313 | 0.033 | 0.466 | 0.265 | 0.174 | 0.000 |
| SE of Mean uHe | 0.037 | 0.029 | 0.041 | 0.037 | 0.019 | 0.022 | 0.035 | 0.043 | 0.000 |

**Key** No. Bands = No. of Different Bands No. Bands Freq. $\geq 5\%$ = No. of Different Bands with a Frequency $\geq 5\%$ No. Private Bands = No. of Bands Unique to a Single Population No. LComm Bands ( $\leq 25\%$ ) = No. of Locally Common Bands (Freq. $\geq 5\%$ ) Found in 25% or Fewer Populations No. LComm Bands ( $\leq 50\%$ ) = No. of Locally Common Bands (Freq. $\geq 5\%$ ) Found in 50% or Fewer Populations He = Expected Heterozygosity = $2 * p * q$ uHe = Unbiased Expected Heterozygosity = $(2N / (2N-1)) * He$ Where for Diploid Binary data and assuming Hardy-Weinberg Equilibrium, $q = (1 - \text{Band Freq.})^{0.5}$ and $p = 1 - q$ .

**Table 5:** Mean Biodiversity Statistics within Populations (Mean±SEM)

| Population | Na | Ne | I | He | uHe |
| --- | --- | --- | --- | --- | --- |
| Abuja | 0.897±0.151 | 1.245±0.057 | 0.215±0.045 | 0.144±0.031 | 0.173±0.037 |
| Adamawa | 0.308±0.117 | 1.106±0.044 | 0.089±0.035 | 0.061±0.024 | 0.073±0.029 |
| Benue | 0.667±0.153 | 1.264±0.065 | 0.206±0.048 | 0.143±0.034 | 0.172±0.041 |
| Kaduna | 1.436±0.141 | 1.444±0.059 | 0.388±0.044 | 0.261±0.031 | 0.313±0.037 |
| Kano | 0.154±0.086 | 1.046±0.028 | 0.042±0.024 | 0.028±0.016 | 0.033±0.019 |
| Borno | 1.949±0.051 | 1.686±0.046 | 0.569±0.022 | 0.389±0.018 | 0.466±0.022 |
| Nasarawa | 1.282±0.156 | 1.355±0.052 | 0.337±0.042 | 0.221±0.029 | 0.265±0.035 |
| Sokoto | 0.692±0.148 | 1.28±0.069 | 0.204±0.05 | 0.145±0.036 | 0.174±0.043 |
| Yobe | 0.487±0.081 | 1±0 | 0 | 0 | 0 |

Na = No. of different alleles; Ne = No. of effective alleles; I = Shannon's Information Index; He = Expected Heterozygosity; uHe = Unbiased Expected Heterozygosity

**Table 6.** Percentage polymorphism of RAPD markers across nine locations in Nigeria.

| Population | Polymorphism (%) |
| --- | --- |
| Abuja | 38.46 |
| Adamawa | 15.38 |
| Benue | 33.33 |
| Kaduna | 69.23 |
| Kano | 7.69 |
| Borno | 97.44 |
| Nasarawa | 64.10 |
| Sokoto | 30.77 |
| Yobe | 0.00 |
| Mean | 39.60 |

Nei’s pairwise genetic distance and genetic identity among nine populations have been presented (Table 7). The Yobe samples were more genetically distant from Benue samples than they were when compared with other locations. In terms of genetic identity, the Sokoto tree were most likely more identical to samples located in Borno State.

**Table 7.** Nei’s pairwise genetic distance (below the diagonal) and genetic identity (above the diagonal) among nine populations.

| Location | Abuja | Adamawa | Benue | Kaduna | Kano | Borno | Nasarawa | Sokoto | Yobe |
| --- | --- | --- | --- | --- | --- | --- | --- | --- | --- |
| Abuja |  | 0.825 | 0.771 | 0.872 | 0.832 | 0.896 | 0.860 | 0.685 | 0.732 |
| Adamawa | 0.192 |  | 0.960 | 0.908 | 0.994 | 0.890 | 0.965 | 0.890 | 0.481 |
| Benue | 0.260 | 0.040 |  | 0.832 | 0.957 | 0.867 | 0.916 | 0.957 | 0.428 |
| Kaduna | 0.137 | 0.097 | 0.184 |  | 0.904 | 0.927 | 0.928 | 0.763 | 0.675 |
| Kano | 0.184 | 0.006 | 0.044 | 0.100 |  | 0.896 | 0.962 | 0.873 | 0.500 |
| Borno | 0.109 | 0.117 | 0.143 | 0.076 | 0.110 |  | 0.924 | 0.978 | 0.754 |
| Nasarawa | 0.151 | 0.036 | 0.088 | 0.075 | 0.039 | 0.080 |  | 0.834 | 0.587 |
| Sokoto | 0.378 | 0.117 | 0.044 | 0.271 | 0.135 | 0.226 | 0.182 |  | 0.337 |
| Yobe | 0.312 | 0.732 | 0.848 | 0.392 | 0.694 | 0.283 | 0.532 | 1.086 |  |

From the Analysis of molecular variance using RAPD markers among locations, it was recorded that estimated variance was more within the populations than among the populations (Table 8). Figure 7a-i shows the test tree at various sampling locations.

**Table 8.** Analysis of molecular variance using RAPD markers among locations.

| Source of variation | Df | SS | MS | Estimated Var. | Variance (%) |
| --- | --- | --- | --- | --- | --- |
| Among populations | 8 | 106.370 | 13.296 | 2.716 | 35 |
| Within populations | 18 | 92.667 | 5.148 | 5.148 | 65 |
| Total | 26 | 199.037 |  | 7.864 | 100 |

Df = degree of freedom; SS = sum of square; MS = mean square; Var. = variance

**Figure 7(a – i):**
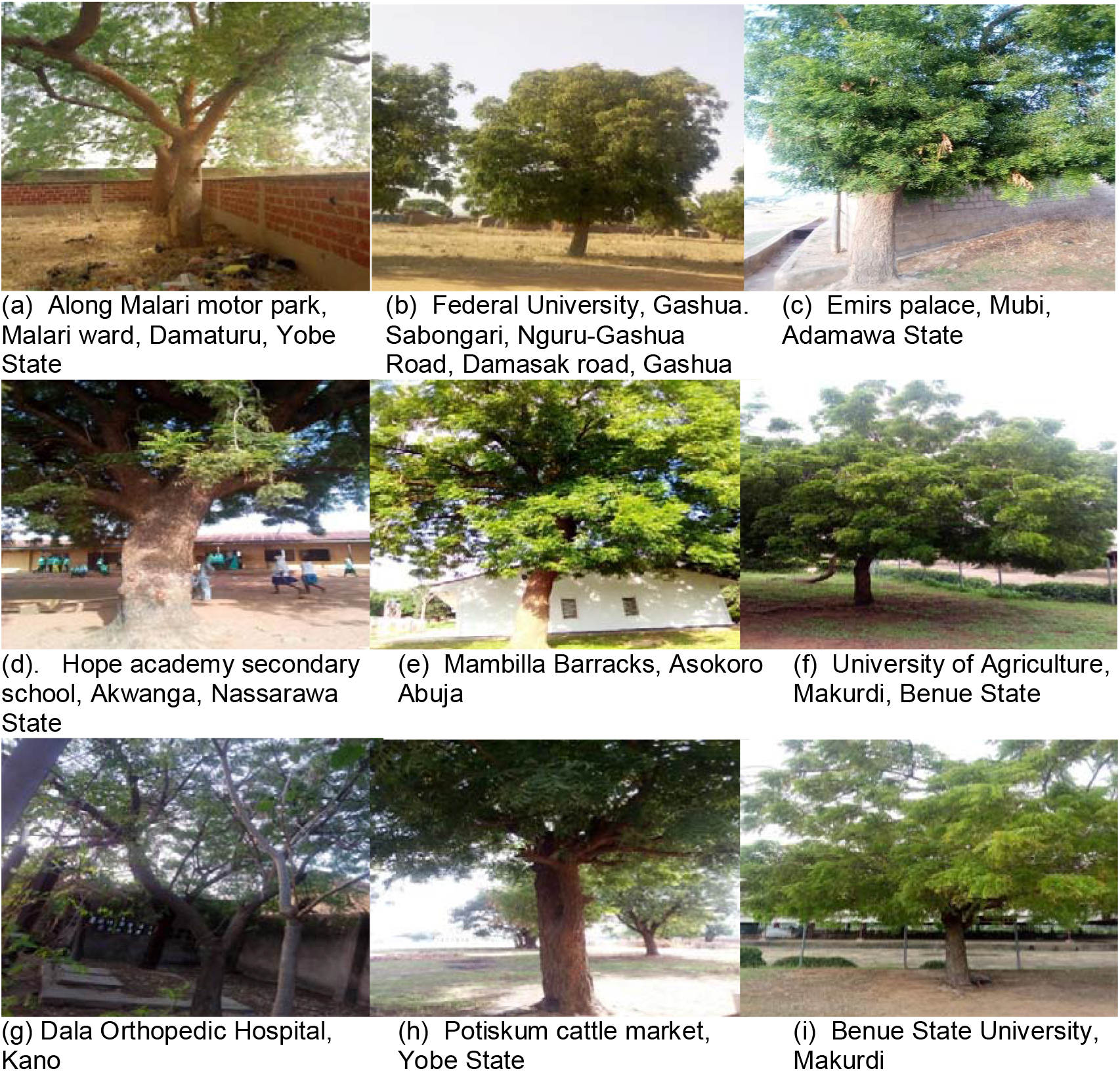
Test tree at various sampling locations. Figure 8 shows Principal Component Analysis of samples. Results showed that Abuja 1 and Abuja 3 samples were distinct from the other groups which formed clusters. The UPGMA Phylogeny of Samples (Figure 9) also showed the district nature of the 2 Abuja tree samples. Among the locations, Adamawa, Nasarawa and Kaduna populations were clustered. This may indicate likely genetic identities (Figure 10). Again, as observed in Figures 4 and 5, the Abuja tree samples were separated from the population (Figure 11) in terms of phylogeny.

## 4.0 Discussion

The findings of this study corroborate previous reports of the neem tree’s economic importance to Nigerians’ socioeconomic lives, especially in the north (see Table 1) (Uko and Kamalu 2001; Lale 2002; Uko and Kamalu 2001; Uko and Kamalu 2001; Uko The tree is also adaptable to a wide range of climatic and topographic conditions, thriving in sandy, stony shallow soil sand as well as soils with hard calcareous or clay pan, according to the report. The neem tree just requires a little water and a lot of sunlight to thrive (Ogbuewu, et al., 2010). The pH differences between soil depths were minor. Organic carbon levels were high (1.61 – 1.97 percent) and magnesium levels were moderate (2.05 – 2.82 Cmole/kg), according to Metson (1961), regardless of location in the North East or soil depth. Similarly, nitrite levels (0.01–0.04 ppm) were similar regardless of position and depth (see Table 5). Iron content, on the other hand, varied, with the lowest concentration in Borno (BN-BK-2) (2.29 – 5.78 ppm). Though iron is classified as a micronutrient since only small amounts are needed for normal plant development, Borno (BN-BK-2) had the lowest iron concentration, ranging from 2.29 to 5.78 ppm. The evaluation of genetic diversity of any species is very important for their conservation and gene manipulation (Khan et al. 2012).

Neem is thought to have a high cross-pollination rate. There have been records of inter-provenance differences in morphological and physiological characteristics in neem. To determine the degree and/or form of genetic (DNA) variation in neem, a powerful molecular technique must be used. The genetic variation of neem ecotypes has recently been determined using molecular techniques such as AFLP, RAPD, ISSR, and RFLP banding patterns (Kota et al. 2006). To extract high-quality genomic DNA, 27 neem samples from Northern Nigeria were used.Nine (09) random primers were obtained and screened for responsiveness to the neem genome, with 5 of them proving to be responsive (Table 2). Because of its ability to search across regions of the genome, RAPD analysis is well suited for phylogenetic studies at the species level. Primers used in RAPD studies are usually random (Kumari and Thakur 2014), necessitating the acquisition of a large number of primers for screening. This increases the likelihood of finding primers that answer. Dhillon et al. (2007) investigated the degree and distribution of genetic diversity in *A. indica* from different eco-geographical regions of the world using RAPD markers and found that they were reliable.

The genetic similarities and variations among the neem samples examined varied significantly from one primer to the next. Variations in RAPD primers were discovered in the study of Neem tree genetic diversity. Primer OPA-15 was the most powerful of the primers used in the analysis. The average number of successful alleles was 1.095–1.625. When compared to the other samples included in the analysis, OPA-15 had the highest diversity index. Heterozygosity was expected to be between 0.088 and 0.405. Primer OPA-15 increased heterozygozity.

Dhillion et al. (2007) used RAPD molecular markers to assess genetic diversity in *Azadirachtaindica*populations from various eco-geographical regions of India. A total of 40 decamer primers were used, and 24 of them resulted in polymorphic banding patterns. A total of 152 distinct DNA bands could be reliably collected, with 104 (68.4%) of them being polymorphic. To classify genetic relationships, polymorphisms were graded and used in band-sharing research. All 36 populations were grouped into two major groups using cluster analysis based on Jaccard’s similarity coefficient and UPGMA.

Using Primer OPA-02, visible bands were reported in samples collected from Abuja-2, Adamawa-3, Benue-1, Benue-2, Borno-3, Sokoto-3 and Yobe-2 (Figure 2). The implication is that among the samples collected from Abuja, one was genetically distinct from the other. This was similar to those collected in Adamawa, Borno, Benue, Sokoto and Yobe States respectively. Findings indicated that samples collected from Abuja were genetically distinct from one another even though they are collected from the same state. This however doesn’t agree with Singh *et al. (*2005) who analyzed the RAPD to access genetic divergence among 29 neem accessions collected from two agro-ecological regions of India (11 agro-climatic sub-zones) and found out that 14 were polymorphic, generating a total of 3 amplifications products with an average of 5.21 products per polymorphic primer and estimated gene diversity of 0.49.

Contrary observation was seen using Primer OPA-03, samples collected from Yobe State were of similar genetic identity as their banding patterns were similar (Figure 4). Figure 5 and 5 also showed majorly dissimilar characteristics where Primer OPA-15 had more than 80% of the samples collected showed amplified bands, with majorly dissimilar characteristics.

Furthermore, a striking result is that they also had similar amplifications with samples collected from Borno State using both OPA-03 and OPA-15 (plates 3and5) indicating possible genetic identities. Therefore the high percentage of similar banding profile generated in this study is not surprising but expected due to the close proximity between the two states. Also, samples from Born and Yobe had broader leaf size and had delayed seed emergence (see Table 18). Farooqui et al. (1998) reported RAPD profiles of 17 accessions of neem from India were generated using 49 random DNA primers. The dendogram of similarities amongst the RAPD profiles suggested that there was less variation than expected within neem from India. In addition, the pattern of RAPD similarities obtained did not correspond to the pattern of geographical variation in neem. This result is not unusual when assessing provenance variation using molecular methods, and the use of additional genetic analyses would have assisted the interpretation of the results.

In the Abuja samples, 20 bands were observed; all these different bands had a Frequency ≥ 5% (Table 5). However, the Borno samples had the highest number of bands (38). Of the 38 bands in the Borno samples, 20 were locally common bands found in 50% or fewer populations.

Indication of significant polymorphism was reported in the study. Percentage polymorphism was highest among the Borno samples (97.44%), compared to samples in Yobe State with no polymorphism. Mean percentage polymorphism was 39.60%. Mean number of available alleles ranged from 0.154 (Kano samples) to 1.949 (Borno samples). In spite of this difference, number of effective alleles between these two locations were comparable (1.046 – 1.686).This is an indication that there is an allelic similarities between Borno and Kano samples. It is most likely that the parent plants were derived from the same mother plant, several years ago. From the Analysis of molecular variance using RAPD markers among locations, it was recorded that estimated variance was more within the populations than among the populations (Table 8). This however does not collaborate with Farooqui et al. (1998) who reported RAPD profiles of 17 accessions of neem from India were generated using 49 random DNA primers. The dendogram of similarities amongst the RAPD profiles suggested that there was less variation than expected within neem from India.

Application of PCA tool and multivariate statistical analysis provide useful means to estimate morphological diversity within and between germplasm collections. The distinctive nature of Abuja samples among other samples (see Fig. 6 and 7) collaborates withMaletsema*et al*. (2019). The genetic relatedness is useful for selection of parental lines for hybridization in crop improvement. Among the locations, Adamawa, Nasarawa and Kaduna populations were clustered. This may indicate likely genetic identities (see Fig. 8). The genetic relatedness is useful for selection of parental lines for hybridization in crop improvement Dosso-Aminon et al. (2015). Abuja tree samples were separated from the population (see Fig. 9) in terms of phylogeny. Maletsema et al. 2019 working on sorghum accessions reported that Accession 10 collected was distantly related with the other accessions which disagrees with the present findings. The diversity observed among the neem samples could be useful in improvement of neem for various traits (Updhyaya et al. 2010).

**Figure 8:**
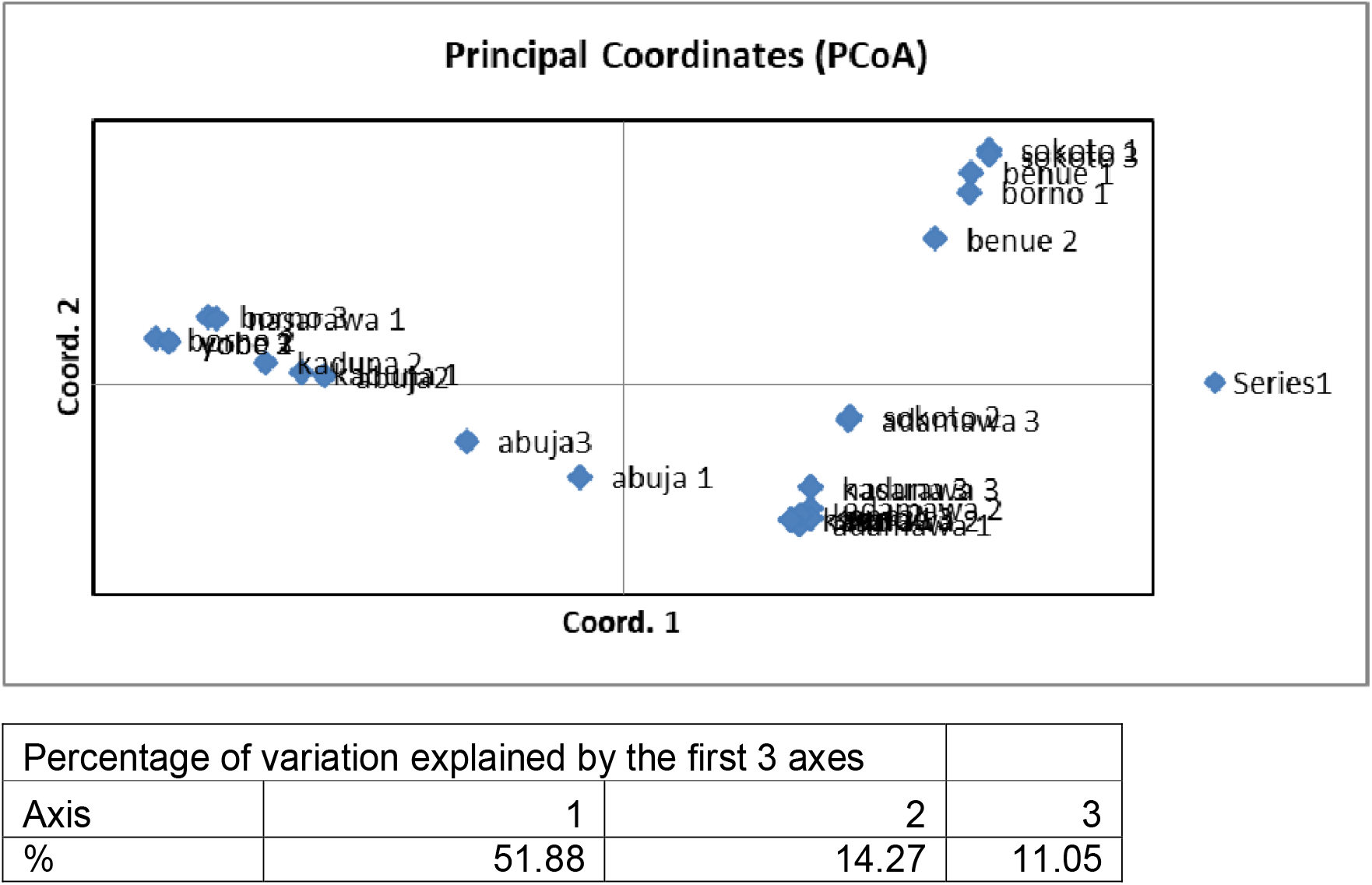
Principal Component Analysis showing Sample Clustering.

**Figure 9:**
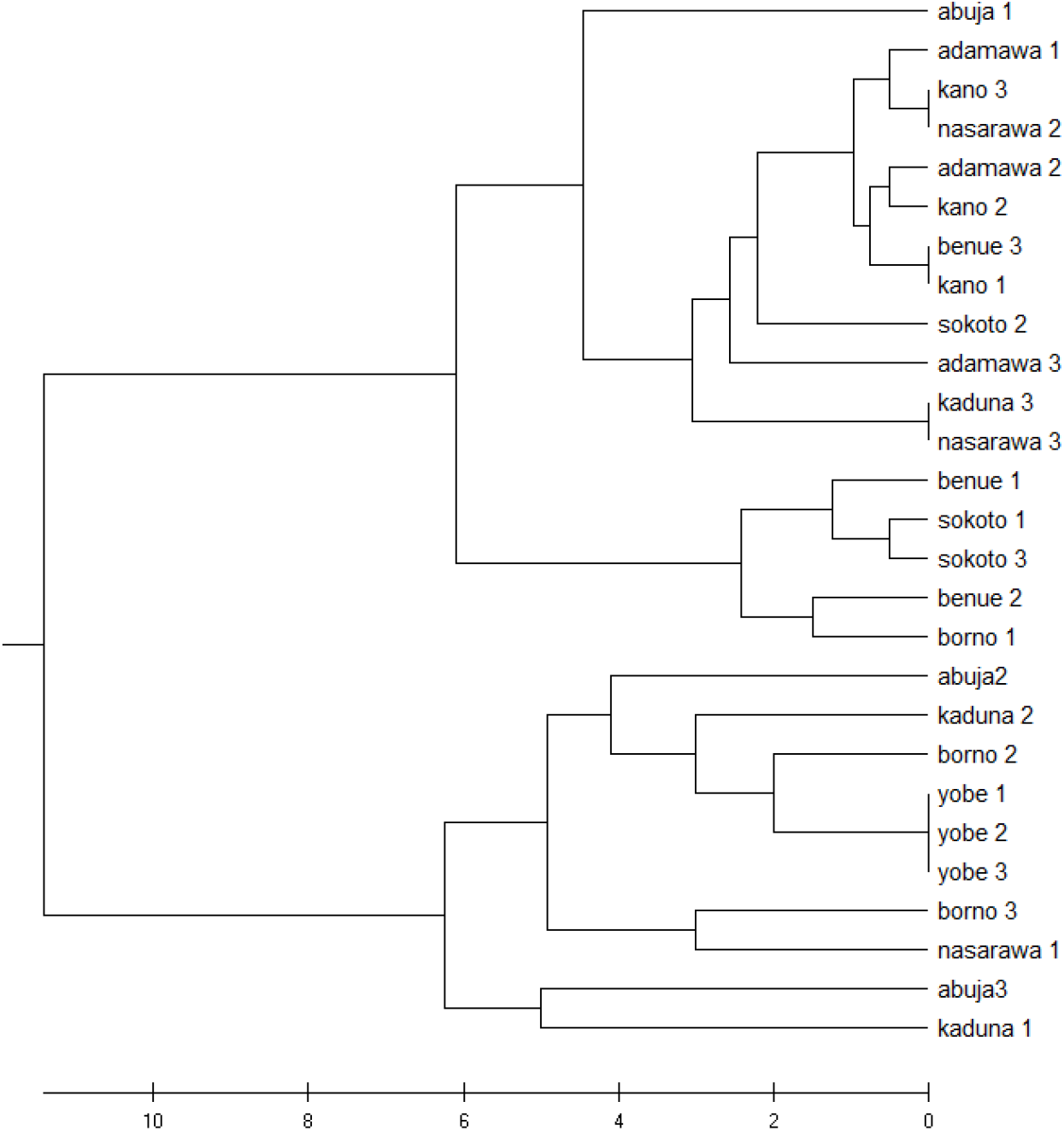
Unweighted pair group method with arithmetic mean phylogeny of samples.

**Figure 10:**
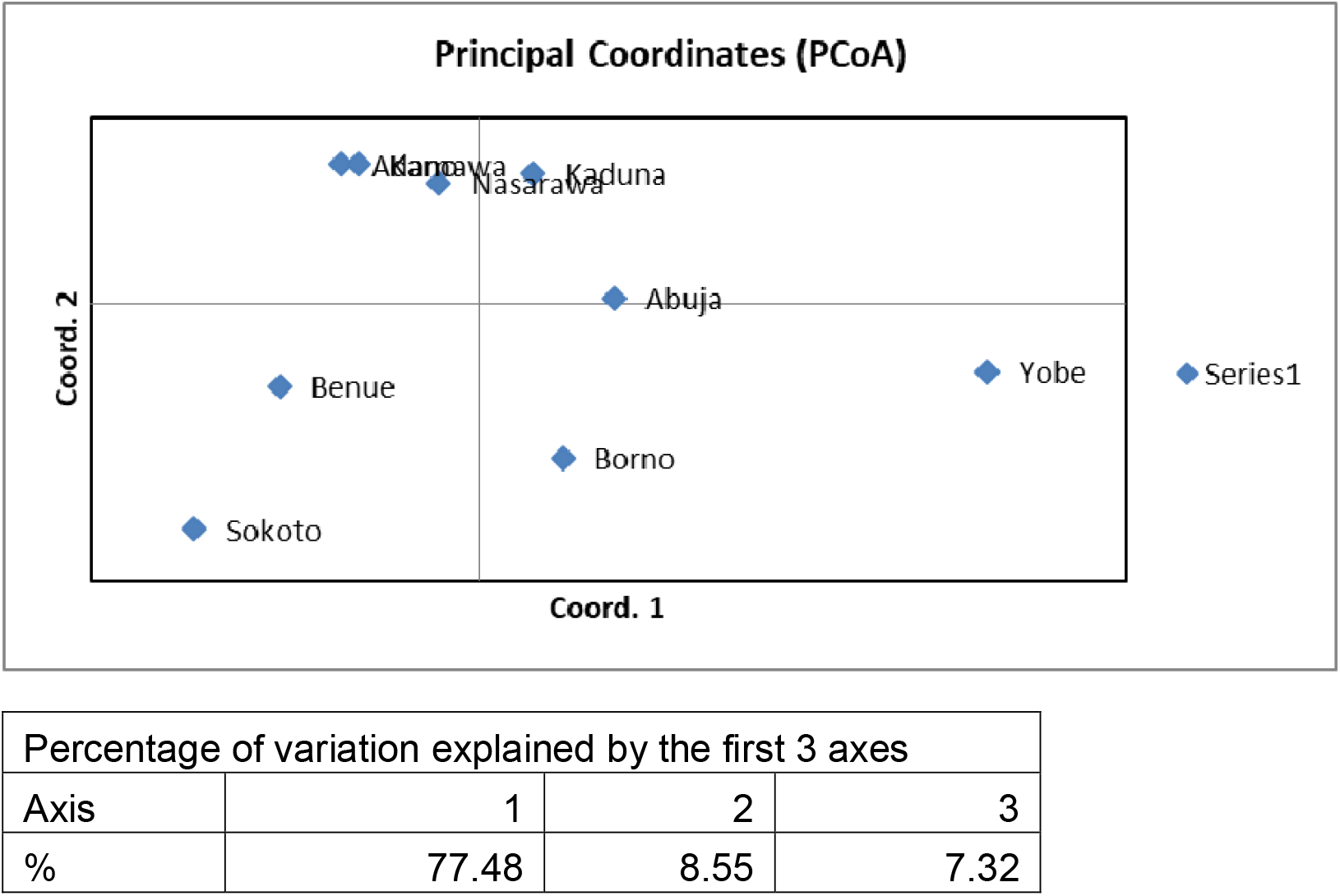
Principal component analyses showing population clustering.

**Figure 11:**
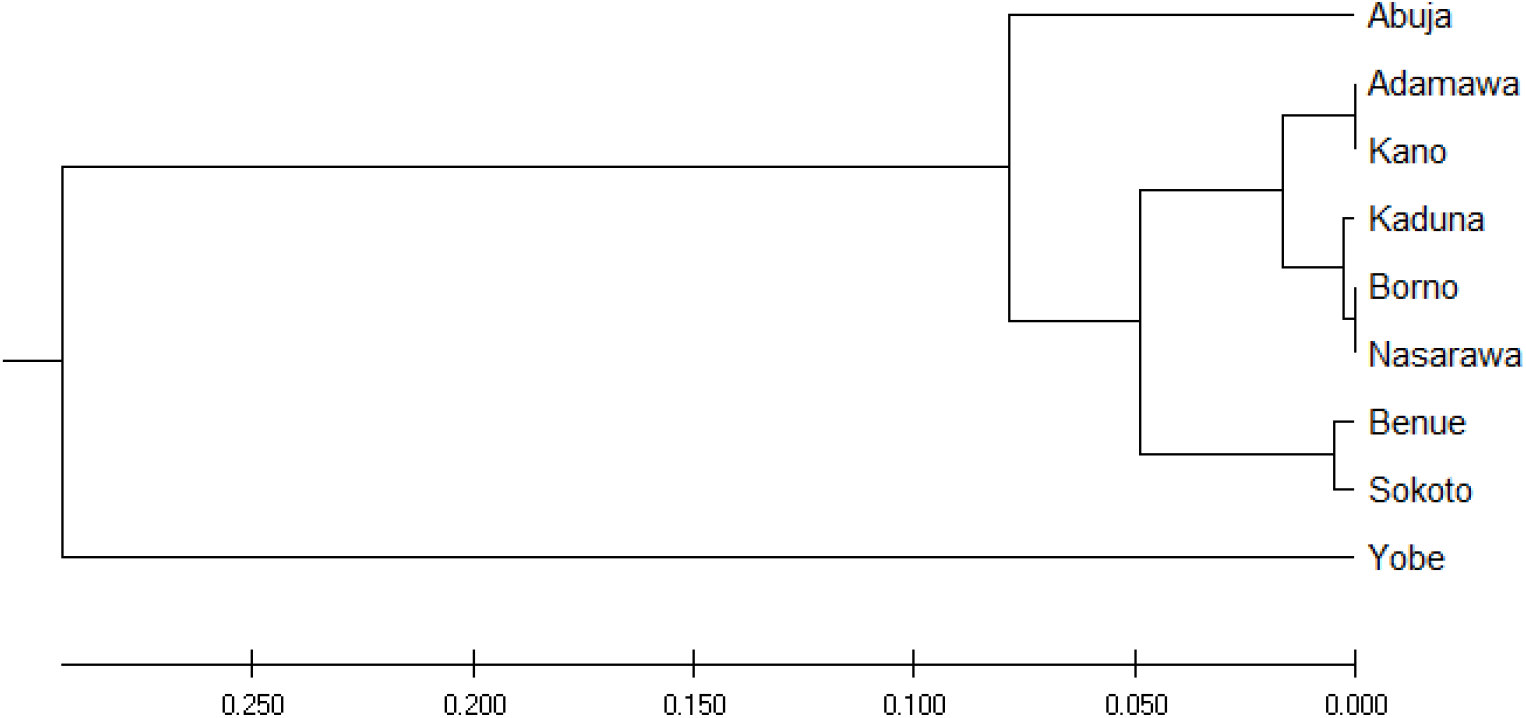
Unweighted pair group method with arithmetic mean phylogeny of populations.

## Conclusion

The study thus showed genetic dissimilarities among the trees investigated. These genetic disparities as observed through RAPD analysesexisted across the ecological zones.It is indeed unclear if this was caused by plant distribution, soil conditions, weather, or atmosphere. These plants, on the other hand, clearly differed in their ecotypes. The recorded potential for genetic diversity is a significant attribute for ecosystem management and germplasm conservation, given the economic value of this tree. Despite the molecular diversity found among neem trees in Northern Nigeria, it had no effect on their economic importance or significance in one area compared to another.

## Authors’ contributions

The study was designed by GOA and BI, and supervised by GOA, BI, and BOE. The field work was carried out by BPM. Data collection and interpretation was by GOA, BI, BOE, and BPM. Statistical analysis of data was by BI. All authors read, revised and approved the final manuscript.

## Acknowledgements

The authors are grateful to the Tertiary Education Trust Fund (TETFund), Nigeria, for providing grants for the study. The assistance and constructive criticisms of Prof. J.K. Mensah of the Department of Botany, Ambrose Alli University, Ekpoma, Nigeria, are highly appreciated.

